# PREDICTION OF SINGLE NUCLEOTIDE VARIANT EFFECTS RELATED TO SEX DETERMINATION IN *Carica papaya (Caricaceae)*

**DOI:** 10.1101/2024.09.15.613137

**Authors:** João Victor Villas Bôas Spelta, Tetsu Sakamoto

## Abstract

Papaya (*Carica papaya*) produces one of the most consumed fruits worldwide, holding great economic importance, especially in tropical regions. Papaya trade mainly involves gynodioecious cultivars with a ratio of 1:1 or 2:1 of hermaphrodites to females. For commercial reasons and also inherent to cultivation, it is preferable to have as many hermaphrodites as possible. In general, it is still not possible to produce only hermaphrodite seeds, so the sex of the plant is usually identified by conventional methods after the first flowering of the papaya plant. This occurs about 4-6 months after planting the seedling, with the females typically being discarded at the end. To avoid wasting resources and achieve higher harvest yields, producers may also resort to molecular methods using sex markers. However, this alternative also has its limitations, including high costs. Given these challenges, many researchers have focused on studying the process of sex determination in *C. papaya*, but the factors directly influencing this process remain unknown. The elucidation of this mechanism is not only of great agronomic interest, but also represents a significant opportunity for *C. papaya* to establish itself as a model organism for studying sexual chromosomes of recent evolutionary origin. Therefore, in this work, a bioinformatics strategy was used to address the topic. A genotype-phenotype association study was conducted to find possible genetic factors involved in sex determination using resequencing data from 36 individuals (24 male papaya plants and 12 hermaphrodites) obtained from public databases. The association study was preceded by a variant calling performed with BCFTOOLS, which found 75,607 variants, leaving 37,027 after filtering. Association studies were then carried out using the PLINK program with the filtered variants, and among these, 251 of the most statistically significant variants were applied to the SnpEff program for variant annotation, returning 449 effects, including 402 with a modifier level of impact, 22 with a low impact, and 25 with a moderate effect. Inferences and gene annotations were also performed using the Augustus software and BLASTP alignments with the gene sequences that had moderate effects predicted by SnpEff, as well as de novo genome assemblies of a male sample and its alignment with the hermaphrodite sex-determining region. These results were recorded and compared with previous studies in the literature. This allowed for the conclusion that the specific results obtained serve as a starting point for more robust studies to understand the molecular mechanisms of sex determination in *C. papaya*.

## 1. Introduction

*Carica papaya*, described by Linnaeus in 1753, is commonly known as papaya tree and stands out as a plant of significant economic importance. The fruit it produces, called papaya, is cultivated in tropical and subtropical regions, with large-scale production primarily in tropical areas (Jiménez; Mora-Newcomer; Gutiérrez-Soto, 2014). In 2022, the main producers were, in ascending order, India, the Dominican Republic, Brazil, Mexico, and Indonesia, with these countries accounting for approximately 10 million tons worldwide (FAOSTAT, 2023). In Brazil, papaya is cultivated in 25 states and the Federal District, with average yields ranging between 4.28 and 5.69 kg per square meter (DANTAS, 2013). In Rio Grande do Norte, the cities of Baraúna, Apodi, and Pureza stand out as important production hubs, significantly contributing to the state’s food production and local job creation (IBGE, 2020).

In addition to being one of the most nutritious and widely consumed fruits globally in its natural form, papaya plays an essential role in the food industry, being processed into various products such as sweets, dehydrated and candied fruits, jams, juices, and pulp (SEBRAE, 2013). The papaya tree is also recognized for its latex, which is rich in papain, a proteolytic enzyme with applications in the pharmaceutical, textile, beverage, leather, and cosmetics industries (Ming; Yu; Moore, 2006). Its nutritional benefits are remarkable, being an exceptional source of vitamins A and C, folate, potassium, niacin, thiamine, iron, riboflavin, calcium, and dietary fiber (Karunamoorthi et al., 2014). The versatility of papaya is evident in its diverse consumption, standing out as a food with unique nutritional value.

The papaya tree is characterized as a fast-growing tropical tree, reaching an average height of 6 meters and rarely branching, producing fruit recurrently over the years, beginning 8–10 months after germination. It is a C3 metabolism plant that produces climacteric fruits in the form of berries, displaying great diversity in size and shape (Jiménez; Mora-Newcomer; Gutiérrez-Soto, 2014). For example, the fruits of hermaphroditic plants tend to be elongated, varying from cylindrical to pear-shaped, while the fruits of female plants tend to be rounded (Chaves-Bedoya; Nunez, 2006). Belonging to the *Caricaceae* family, which includes *C. papaya*, this family comprises 35 species distributed across six genera, predominantly dioecious, with male and female flowers on separate individuals. However, there are exceptions, such as the species *Vasconcellea monoica* and *V. cundinamarcensis*, which are monoecious, presenting both male and female flowers on the same plant. *C. papaya*, however, is one of only two trioecious species in its family, presenting hermaphroditic, female, and male individuals (Ueno *et al*., 2014). In agricultural production, the most common cultivars are predominantly gynodioecious (Valverde *et al*., 2019). On the other hand, wild populations of *C. papaya* are predominantly dioecious, with staminate and pistillate flowers on separate individuals (Chae; Harkess; Moore, 2021).

About 90% of flowering plant species are hermaphrodite, an advantageous characteristic for ensuring reproduction, as it allows, in most cases, self-fertilization. Besides, only 6% of all angiosperms are dioecious, presenting only staminate (male) or pistillate (female) flowers on each individual (Carvalho; Renner, 2014). The distribution of sex alleles in *Carica papaya* reveals an intriguing evolutionary scenario, reflecting the complexity of sexual strategies in plants. The sex chromosomes present in *C. papaya* can be X, Y, or Yh, following the XY pattern. Regarding genotypes, they can be: xx, the female being homozygous recessive, with mm alleles; xY (male) and xYh (hermaphrodite), heterozygous, with alleles M1m and M2m, respectively (Hofmeyr, 1938). These alleles play a crucial role in sex determination in *C. papaya*, and their crosses are more clearly demonstrated below (table 1).

**Table 1.** **Allelic combinations and possible genotypes in *C. papaya***

| Alleles | M1 | M2 | m |
| --- | --- | --- | --- |
| M1 | M1M1* | M1M2* | M1m (male) |
| M2 | M1M2* | M2M2* | M2m (hermaphrodite) |
| m | M1m (male) | M2m (hermaphrodite) | mm (female) |

\*Lethal genotypes in red.

It is important to note that all combinations that do not carry the recessive female allele are lethal, indicating a possible degeneration of genes on the Y and Yh chromosomes due to the accumulation of mutations in these chromosomes and the absence of genes necessary for complete embryonic development, both in the Y and Yh (WANG *et al*., 2012).

When discussing the evolutionary history of chromosomes and the origin of the species itself, some aspects stand out. In the literature, the genetic similarity between the Y and Yh chromosomes is well established, ranging from 98.9% to 99.6%. The greatest divergences occur in the intergenic and repetitive regions, revealing that these chromosomes are much more closely related to each other than to the X chromosome (Yu *et al*., 2008). It is estimated that the Y and Yh chromosomes separated relatively recently on the evolutionary scale compared to the X chromosome, with phylogenetic studies indicating that X and Y diverged 0.5 to 2.5 million years ago. It can be said that the sex chromosomes in *C. papaya* are of recent origin, reaffirming their importance as a model for studying primitive sex chromosomes (Ming *et al*., 2008).

Although the origin of the species lacks specific archaeological evidence, its presence prior to the arrival of Europeans and the analysis of wild haplotypes suggest a Mesoamerican origin, more precisely in territories that now belong to countries such as Belize, Guatemala, and southern Mexico (Vanburen *et al*., 2015). Additionally, phylogenetic analyses reveal dioecism as the norm in the Caricaceae family, with *C. papaya* having diverged over 27.5 million years ago from the most closely related species, *Vasconcellea monoica* (Xavier *et al.,* 2021). It is hypothesized that the hermaphroditic trait, in the form of the Yh chromosome, re-emerged evolutionarily approximately 4,000 years ago, coinciding with the emergence of agriculture in this region of Central America. Furthermore, the lower diversity of Yh strongly indicates a genetic bottleneck resulting from a domestication process, which may have occurred in a pre-Columbian context by civilizations in the region, such as the Maya. Therefore, it is understood that hermaphroditic *C. papaya* plants evolved from male plants, possibly selected by humans due to their favorable phenotype (Vanburen *et al*., 2015).

Considering that the hermaphroditic trait is characterized as the evolutionary base in angiosperms, but also that the *Caricaceae* family is almost entirely composed of dioecious species, as well as the wild *C. papaya* specimens, it is possible to estimate the steps that may have led from hermaphroditism to dioecy in wild papayas. This transition to dioecy in plants involves the emergence of a genetic regulatory mechanism through sex chromosomes, which may involve intermediate pathways such as androdioecy or gynodioecy (Zerpa-Catanho *et al*., 2019). However, as previously mentioned, hermaphroditism in *C. papaya* does not appear to be a result of an ancestral trait but rather of a domestication process, which adds complexity to the attempt to unravel its sexual differentiation mechanism (Vanburen *et al*., 2015).

One of the most accepted models for the transition from hermaphroditism to dioecy involves two genes responsible for the transition, where two mutations in separate loci are thought to have occurred. In male heterogametic systems (XY), such as in *C. papaya*, it is estimated that an initial mutation, likely recessive, resulted in male sterility and the formation of gynodioecious populations (females and hermaphrodites). The second mutation, which was eventually dominant, would induce female sterility, leading to a dioecious population with separate sexes. These loci would have remained linked in a sexual determination region, with sex chromosomes consolidated through complete disruption of recombination via repetitive regions and inversions incorporated into the chromosomes. Additionally, it is understood that an individual with both mutations would be completely sterile, which would facilitate the cessation of recombination between the primitive chromosomes (Charlesworth, 2002). Concurrently, support for this two-gene model is found in other plant species that also utilize sex chromosomes, such as asparagus (*Asparagus sp*), Silene latifolia, kiwi (*Actinidia deliciosa*), and date palm (*Phoenix dactylifera*). The adaptive advantage that dioecy provides is that by inhibiting self-fertilization of flowers, it confers greater genetic variability to populations (Zerpa-Catanho *et al*., 2019).

When characterizing the genome and sex chromosomes of *C. papaya*, it is noteworthy that the species has a relatively small genome compared to *Arabidopsis*, with 351.5 Mbp and a distribution of 2n = 18, with homomorphic chromosomes of metacentric and submetacentric conformations, comprising 63.49% adenine/thymine and 36.51% guanine/cytosine, and with the sizes of the 9 autosomes being virtually identical (Vanburen *et al*., 2015). Despite the genomic complexity presented by the sex chromosomes of *C. papaya*, linked to redundant sequences and the presence of retrotransposons, many authors have focused on describing the sex-determining regions among the X, Y, and Yh chromosomes. Present in the latter two are the regions known as MSY (Male Specific Region) and HSY (Hermaphrodite Specific Region), respectively. These two regions are located near the centromere of their chromosomes and are approximately 8 - 9 Mbp in size, corresponding to 10% of the chromosomes (Chae; Harkess; Moore, 2021). In contrast, the sex-determining region of the X chromosome is only 3.5 Mbp. The difference in size is attributed to the presence of large retrotransposons (Aryal; Ming, 2013), with two significant inversions standing out. Chronologically, it is understood that the first inversion caused the suppression of recombination between X and the Y chromosomes 2-3 million years ago, while the second inversion was capable of expanding it (Wang *et al*., 2012).

In HSY, the number of transcription units found was 94, including 66 protein-coding regions and 28 pseudogenes. Both HSY and MSY are surrounded by large pseudoautosomal regions (Chae; Harkess; Moore, 2021). Combined, the two regions were described with 14,528 nucleotide substitutions and 965 indels from a direct comparison between HSY and MSY by Ueno *et al*. (2014). This comparison also revealed the presence of 294 unmapped regions containing gaps, as well as 14 putative genes with polymorphisms (Ueno *et al*., 2014). It is worth noting that the high genetic similarity between the Y and Yh chromosomes makes it difficult to identify genetic differences between the two, mainly because most differences are located in introns rather than exons, and also due to the presence of large methylated regions (Aryal *et al*., 2014; Zerpa-Catanho, 2019).

Regarding the HSY region and consequently the hermaphroditic phenotype, they have commercial significance for at least two reasons listed in the literature. Firstly, because the current commercialization of papayas is predominantly done with gynodioecious cultivars and hermaphroditic fruits. The first reason is that fruits from hermaphroditic plants have a pear-shaped, more elongated form, whereas fruits from female plants are more rounded with a larger central cavity, making them less preferred by markets that distribute the fruits to consumers (Xavier *et al*., 2021). The other reason for preferring hermaphroditic papaya plants relates to cultivation practices, where female papaya plants are naturally incapable of self-fertilization and require the planting of male or hermaphroditic individuals nearby to produce fruit (Aryal; Ming, 2013). This situation is further exacerbated for producers using dioecious cultivars, who may lose up to 10% of their planting space to plants that do not produce fruit. However, most farmers using gynodioecious cultivars also face similar issues. Due to heterozygosity, crosses between hermaphroditic plants can produce seeds that are both hermaphroditic and female in a 2:1 ratio, and crosses between female and hermaphroditic plants yield a 1:1 ratio (Ming; Yu; Moore, 2006).

Therefore, a considerable portion of papaya plants are discarded 3 to 5 months after germination because they are female (Ming; Yu; Moore, 2006). This occurs with the identification of sex based on the observation of the first flowering, near the time of planting the seedlings. Typically, producers plant three or four seedlings in the same planting bed to achieve up to a 90% chance of having a hermaphrodite. The discarding of these female seedlings represents a waste of resources such as water, agricultural inputs, as well as labor time and available planting space (Honoré *et al*., 2020).

A better understanding of the sexual determination process in *C. papaya*, as well as the identification of the genetic factors involved in this process, would advance methodologies aimed at cultivating only hermaphrodites, in order to avoid the waste of resources from discarding female plants. Additionally, it presents an ideal opportunity to understand how recent sex chromosomes on the evolutionary scale originate and differentiate. It is also worth highlighting C. papaya as an excellent model for studying sexual determination in plants that also use sex chromosomes, due to its advantages such as rapid growth, low maintenance, a relatively small and already sequenced genome, and the ability to be cultivated both in vitro and in vivo (Aryal *et al*., 2014).

Throughout the 20th and 21st centuries, many efforts have been made to solve the mystery of sexual determination in *C. papaya* and to address the challenges of its cultivation. This dissertation will present some of these efforts to provide the reader with an understanding of the current landscape and will then demonstrate the present attempt to deepen the knowledge of sexual determination.

The identification of sex through molecular techniques has emerged as a way to mitigate the situation regarding resource wastage. Numerous researchers have developed sex markers that enable this identification, usually done with seedlings or seeds before germination. These markers include RFLP (Restriction Fragment Length Polymorphism), RAPD (Random Amplified Polymorphic DNA), AFLP (Amplified Fragment Length Polymorphism), SNP (Single-Nucleotide Polymorphisms), microsatellites, and microarrays (Avila-Hernandez *et al*., 2023).

One of the earliest markers was developed by Urasaki *et al* using RAPD in 2002. It had 225 base pairs and allowed the identification of the sex of *C. papaya* seedlings between hermaphrodites and males. However, due to producing a higher number of false negatives and some inconclusive results, the authors opted to use the papain gene as a control along with two other markers to mitigate these issues (Urasaki *et al*., 2002).

Additionally, in 2002, Deputy *et al*. produced oligonucleotides that can detect sex in *C. papaya* both in gynodioecious and dioecious cultivars with a prediction accuracy of 99.2%. These primers, named W11 and T12, are made from papaya leaves but can also detect only males and hermaphrodites. To identify females, their team used primer T1, which can amplify sequences from all three chromosomes and is typically used in conjunction with the other two primers to achieve the cited accuracy.

Other notable markers include sex-specific RAPD markers, such as those demonstrated by Chaves-Bedoya and Nunez (2006), and those using other techniques like fluorescence in situ hybridization (FISH) (Abreu; Carvalho; Soares, 2015) and loop-mediated isothermal amplification (LAMP) (Hsu; Gwo; Lin, 2012). The latter allows on-site testing using leaf material, providing results in less than an hour through a color change in the instrument. However, even with methods like this, the use of markers for sexing still requires a high cost for large-scale application, along with specialized labor and equipment (Honoré *et al*., 2020). Other drawbacks include variability in primer specificity depending on the cultivar and, notably, the fact that about 60% of seeds do not germinate after sexing due to the DNA extraction process, making seedling sexing more viable than seed sexing. Nonetheless, the inherent benefits of molecular techniques in terms of water, fertilizer, time, and space savings for producers are clear, as they do not need to wait six months for manual sexing or spend resources on plants that will not be commercialized (Xavier *et al*., 2021).

In the context of sexual determination in *Carica papaya*, literature searches reveal a remarkable variety of approaches undertaken to uncover the genetic factors underlying this crucial process. Throughout the investigations, it becomes evident that sexual determination is possibly influenced by a range of distinct aspects, including phytohormonal effects, epigenetic controls, environmental influences, factors related to the type of sex chromosome, candidate sex-determining genes, and gene regulations (Avila-Hernandez *et al*., 2023). These topics have garnered considerable attention due to the complexity involved in determining the sex of this plant species, prompting a wide range of hypotheses and experiments aimed at unraveling the mechanisms involved in the process.

It is estimated that phytohormones may influence sexual determination in *Carica papaya* directly through flower development. These inferences are based on studies that have evaluated the hormonal regulation of sexual organ development in various plant species. In papaya, genes related to the biosynthesis and signaling pathways of phytohormones, such as ethylene, abscisic acid, auxin, cytokinin, and gibberellins, have been identified in the genome, with differential expression between sexes (Ming *et al*., 2012). In other species, such as melon (*Cucumis melo L.*), the gene *CmACS-7*, involved in the ethylene biosynthesis pathway, suppresses stamen development in female plants, while in male plants, it is suppressed by the gene *CmWIP1*, allowing normal stamen development (Boualem *et al*., 2008) (Martin *et al*., 2009). In grape (*Vitis vinifera*), an allele of a cytokinin metabolism gene is theorized to be an important factor in sexual determination (Fechter *et al*., 2012). For *Zea mays*, the phytohormones brassinosteroid and jasmonate suppress the development of the maize tassel (Acosta *et al*., 2009).

Some researchers have also utilized exogenous applications of phytohormones or their inhibitors and observed effectiveness in altering the sexual expression of plants. For example, literature reports treatments with gibberellin inhibitors in male *C. papaya plants*, which led to the induction of carpel development, highlighting the role of gibberellins in maintaining male characteristics (Kumar & Jaiswal, 1984).

Studies have also demonstrated that auxin concentration is crucial for gynoecium development, with inhibitors of polar auxin transport such as NPA (N-1-Naphthylphthalamic acid) restoring female fertility and resulting in rudimentary hermaphroditic flowers (Zhou *et al*., 2019). Thus, phytohormones emerge as significant factors in sexual determination in *Carica papaya*, interacting in a complex manner to regulate floral development and sexual expression.

Sexual determination in plants is a complex phenomenon involving interactions between genetic, epigenetic, and environmental factors. Epigenetics, in particular, plays a crucial role in promoting sexual phenotypes in plants through modifications to histones, such as methylation or acetylation, and the action of microRNAs that regulate floral development and adaptation to adverse environments (Lin *et al*., 2016; Latrasse *et al*., 2017; Li *et al*., 2021).

Recent studies have focused on uncovering the epigenetic mechanisms involved. An analysis of small RNAs (sRNA) from male, female, and hermaphroditic papaya flowers revealed fourteen microRNA (miRNA) sequences that are differentially expressed among the three sexes. miR169 was found to be most expressed in male flowers, repressing floral development genes and, together with miR160, miR167a, miR167b, and miR393, regulating the auxin signaling pathway, which is essential for carpel and gynoecium development (Aryal *et al*., 2014; Zhou *et al*., 2019).

The phenomenon of sexual reversal, which occurs in some papaya plants, is crucial when discussing epigenetics and environmental factors. Sexual reversal is a critical process of changing sexual characteristics and fruit malformations due to genetic, environmental, and epigenetic factors, or even chemically induced in hermaphroditic and male plants, depending on their genotype. It can occur from hermaphroditic to male or vice versa, resulting in no fruit production and economic losses (Ramos *et al*., 2011; Lin *et al*., 2016; Liao, Yu, Ming, 2017). Lin *et al*. (2016) reported epigenetic marks in the sexual reversal from male to hermaphroditic induced by low temperature, such as methylation of lysine 9 on histone H3 (H3-K9), microRNAs, and hormone-related genes, mainly auxins involved in gynoecium development, which were positively regulated in the reversal, showing more similarity to hermaphroditic than male flowers, thus highlighting the role of epigenetic modification in suppressing the gynoecium due to temperature change. Therefore, sexual reversal adds complexity to understanding sexual determination in *C. papaya* while providing clues that are still being elucidated.

As research into epigenetic regulation in papaya continues, other species are also being studied, allowing comparisons with *C. papaya*. For example, sexual determination in maize is regulated by several non-coding small RNAs through DNA methylation (Parkinson, Gross, Hollick, 2007; Hultquist, Dorweiler, 2008). In the dioecious species *S. latifolia*, cytosine methylation of DNA is necessary to maintain the unisexual nature of male plants. Demethylation of cytosine residues by 5-azacytidine (a cytidine analogue that inhibits DNA methylation) reverts male flowers to hermaphroditic (Janouek, Sirok, Vyskot, 1996).

Another example is the epigenetic modification of the *CmWIP1* locus in melon, caused by the insertion of a transposable element, leading to the development of a unisexual female flower in melon (Martin *et al*., 2009). In *Populus trichocarpa*, the sex-determining region of chromosome 19 has hotspots of small RNAs, suggesting a possible epigenetic mechanism in sexual determination in this species (Tuskan *et al*., 2012). It is also important to note the role of microRNAs, which are hypothesized to play a role in the evolution of unisexual flowers in cucumber (*Cucumis sativus*) by aborting carpel development under environmental stress episodes (Sun *et al*., 2010).

Although efforts to identify the genes responsible for sexual determination in *Carica papaya* have been challenging, new research continues to advance scientific knowledge year after year. These studies not only identify potential genes involved in sexual determination but also pave the way for confirming their roles.

One of the main candidates for a sex-determining gene in *C. papaya* is *Cp2671*, also known as the *SVP* (Short Vegetative Phase) homolog. This gene was identified through transcriptome analysis and high-throughput sequencing (Ht-SuperSAGE) and shares 85% sequence identity with the similar gene in *Arabidopsis thaliana*, where *SVP* is crucial for processes such as flowering regulation, the shift from vegetative to reproductive growth, and the regulation of B and C homeotic genes, which are important for proper floral organ development (Lee *et al*., 2018; Chae, Harkess, Moore, 2021). In *C. papaya*, *SVP* is expressed only in male and hermaphrodite flowers, suggesting it could serve as a sexual marker and might play a role in suppressing carpel development (Ueno *et al*., 2014).

Evidence points to *SVP* producing divergent protein products between Y and Yh chromosomes. The *SVP* protein present in the MSY is functional; however, in the HSY allele, a retrotransposon insertion causes the resulting protein to have an inactive conformation, containing only the K-box domain but lacking the MADS-box domain (Chae, Harkess, Moore, 2021; Ueno *et al*., 2014). This gene stands out as the only one among 14 others with differential expression between hermaphroditic and male flowers, showing a significant increase in expression both at the early and final stages of flower development (Chae, Harkess, Moore, 2021).

Other potential candidates with less prominence include *CpCAF1AL* and *CpSERK*, which possess sex-specific polymorphisms identified by Lee and colleagues in 2018. *SERK* genes (Somatic Embryogenesis Receptor Kinase) are involved in plant growth and development, regulating processes like male sporogenesis, floral organ separation, and embryo development (Lee *et al*., 2018).

Zerpa-Catanho *et al*. (2019) also identified another potential candidate for sexual determination alongside the *SVP* homolog: the *Cp12204* gene, which encodes monodehydroascorbate reductase (MDAR). However, it is believed to have less involvement in sexual determination since this gene is found on all three sex chromosomes, intact on the X and dysfunctional on the Y and Yh chromosomes, without distinguishing them. Furthermore, since MDAR is presumed to be involved in the removal of reactive oxygen species in cellular metabolism, it is thought to be more related to the lethality of genotypes lacking the X chromosome (Ueno *et al*., 2014).

The sexual determination in *Carica papaya* remains a complex and intriguing phenomenon that has not yet been fully elucidated. Understanding this process not only opens doors to cultivating higher yields of hermaphrodite fruits, which are commercially favored, but also offers a unique opportunity to use *Carica papaya* as a model for studying other species with recently evolved sex chromosomes. In this work, it was hypothesized that genetic factors on the Y and Yh chromosomes play a crucial role in sexual determination in *Carica papaya*. These factors are believed to include a gene suppressing carpel development, which is inactive in hermaphroditic papayas but responsible for inhibiting female reproductive systems in male plants. However, further in-depth studies are necessary to confirm this hypothesis and fully understand sexual determination in this species.

In Brazil, where papaya is a significant crop, these findings could have important implications for improving hermaphrodite fruit production, which dominates the commercial market due to its more favorable shape and higher market value. Advances in genetic understanding could lead to breakthroughs in papaya breeding programs, potentially boosting the country’s papaya industry and reducing the economic impact of sex-related fruit production issues.

## 2. Methods

The DNA sequencing data for the Y and Yh chromosomes were obtained from public databases, specifically NCBI (National Center for Biotechnology Information). In total, data from 36 resequenced *C. papaya* samples were obtained, produced in experiments by VanBuren *et al*. in 2015, where 24 individuals were wild and dioecious cultivars and the remaining 12 were hermaphroditic and gynodioecious cultivars. The length in base pairs of the resequenced chromosomes was 8.1 mb. The resequencing data were obtained in FASTQ format using the fastq-dump package, which is part of the SRA-Toolkit (Durbrow, 2018). A FASTQ file is a text file format used to store sequences, in this case, DNA nucleotide sequences along with associated quality information.

The reference genome was also obtained from NCBI, from a resequencing done by Fujian Agriculture and Forestry University, China, in 2022, which used only sequences related to the HSY region. After obtaining the data from the 36 samples and the reference genome through commands on the remote server, the genome was indexed for further steps using the BWA (Burrows-Wheeler Aligner) software (Li; Durbin, 2009).

Quality control of the data was carried out using the Trim Galore tool (Krueger, 2021). It works by incorporating functions from Cutadapt and FastQC to perform tasks such as detecting and removing adapters from sequences, analyzing the quality of sequencing reads, and removing those that are not suitable. The standard package parameters were used, including a Phred score of 20, a maximum error rate of 0.1 per sequence length, and a minimum sequence length of 20 base pairs, among others.

The trimmed FASTQ files were then aligned with the reference genome using the mem function of the BWA software. This command performs alignment based on small sequences with maximum equivalence between the sequences and the reference genome according to the BWA-MEM algorithm. These sequences are then extended using the Smith-Waterman algorithm (Smith; Waterman, 1981). The alignment parameters were the default ones for the package, and the command used generated output files in SAM (Sequence Alignment Map) format.

The SAM files for each sample were converted to BAM (Binary Alignment Map) files, which are equivalent to SAM but in binary format. This conversion facilitates data compression and indexing, and increases processing speed. Additionally, the data within the BAM files were sorted. Both processes were performed using the SamTools software (Danecek *et al*., 2021).

Subsequently, the bcftools program was used to convert the BAM files to BCF (Binary Call Format) and then to VCF (Variant Call Format), enabling variant calling. Both formats are text files used to store genetic variant information and follow the same logic as SAM/BAM for data storage. Once the VCF files for all 36 samples were generated, they were compressed, indexed, and merged into a single VCF file using the VCFTools package (Danecek *et al*., 2011).

The combined VCF file then underwent filtering steps to retain only variants that met the criteria for the next analyses. During this filtering, variants were removed based on the following criteria: (1) QUAL score less than 25; (2) coverage less than 10 reads; (3) more than 50% missing data; (4) minor allele count less than 3; and (5) allele frequency less than 0.05. These procedures were assisted by the VCFTools and BCFTools programs. Additionally, complementary filters were applied using the varFilter package from the vcfutils.pl program to ensure the removal of inadequate variants. In total, the filtering occurred in two stages: an initial phase with only filters (1) and (2) applied to individual samples, followed by a second filter when all samples were combined into a single VCF, using the remaining criteria as well as varFilter.

To perform the association analysis, the PLINK tool (Purcell *et al*., 2004) was used. For this, the VCF file containing the combined data from all samples was converted into .map and .ped format files. The .map file contains genomic variant identifiers, their respective chromosomes, and positions in base pairs. The .ped file is used for pedigree distinction, containing the respective statuses and genotypes of the samples (Chang, 2022). This file was edited to identify the phenotypes of each sample, with the number 1 representing male sequencings and the number 2 representing hermaphrodites. Using the PLINK software, the PED (Pedigree File) was converted to the binary BED (Binary PED File) format, and the association analysis was conducted. The results were visualized through a Manhattan plot, produced by the statistical program R using the qqman library (Turner, 2018).

The choice of PLINK for this analysis was based on its ability to statistically test large quantities of genomic data to identify significant associations between genotypes and phenotypes. The process involves analyzing patterns in allele frequency in genetic variants of individuals that are phylogenetically close but phenotypically distinct (Uffelmann *et al*., 2021). This method can reveal single nucleotide variants associated with the presence or absence of a phenotype. The choice of this method necessitates the completion of the previous steps to ensure the correct input data for PLINK.

After obtaining the results from PLINK, variants with a p-value less than 5 x 10^-8 were selected based on indexed positions. Concurrently, the reference genome (HSY) was subjected to gene prediction using the Augustus software (Stanke, 2018). By submitting the files generated in Augustus to SnpEff along with the reference genome, a *Carica Papaya* database was manually created. This was necessary due to the absence of the species in databases already covered by the native databases of the program. Subsequently, the selected variants were submitted to SnpEff for effect prediction. SnpEff is a bioinformatics tool used to analyze and annotate genetic variants in DNA sequences. It is particularly useful for identifying and characterizing the functional consequences of genetic mutations, such as those statistically associated with sexual differentiation between male and hermaphrodite papaya samples (Bohry, 2021).

As previously mentioned, the annotation of the HSY region was performed using the Augustus software to manually create the database in SnpEff. Augustus is programmed for gene prediction using Hidden Markov Models (HMMs) to identify gene coding locations in DNA sequences. This process provided several important files for proceeding with SnpEff analysis, including GTF (Gene Transfer Format) and GFF (General Feature Format) files, as well as files containing amino acid sequences, coding sequences, exons, and a summarization file.

Subsequently, using the genes predicted by Augustus and the variant annotations in SnpEff, BLASTPs (Basic Local Alignment Search Tool Protein) were conducted with variants identified as having moderate effects in SnpEff. The choice of these variants is due to the fact that, although not classified as highly impactful, moderate effect variants can significantly alter protein function, which may influence important biological processes such as sexual determination.

The purpose of the BLASTP was to identify homologous proteins altered by these variants and compare them with proteins already known in other organisms. This is particularly useful for investigating whether the variants affect genes or genomic regions associated with sexual differentiation, such as transcription factors, genes involved in hormonal pathways, or regulatory elements. Comparing mutated proteins with databases can reveal similarities with genes previously described in other sexual determination systems or related species. This approach may provide insights into which genes or pathways are critical for sexual differentiation in *C. papaya*, contributing to a better understanding of the molecular factors responsible for this process.

The de novo genome assembly was performed using a male *C. papaya* sample, selected from the 24 individuals mentioned previously, based on the highest coverage and total number of reads. This sample, with the access number ’SRR1822135’ in the SRA database, has 10.6G bases. Since the sequence had already undergone trimming, it proceeded to genome assembly, which employed the SPAdes software, widely used for genome assembly from next-generation sequencing data. The assembly process generated a set of scaffolds, which were then compared to the hermaphroditic reference genome using an alignment with D-genies, producing a dot plot graph for visualization. The arrangement of data in a dot plot facilitates the analysis of similarity between the assembled genome and the reference genome, as well as the assessment of assembly quality.

## 3. Results and Discussion

### 3.1 Variant Calling

The present research performed variant calling based on genomic data from Bioproject: PRJNA271489, deposited in NCBI. After performing variant calling following the alignment of the HSY region and the 36 samples, 69,227 single nucleotide variants (SNVs) and 6,380 indels were identified. After the second filtering of the VCF file described previously, 35,361 SNVs and 1,666 indels remained. The results are summarized below (Table 2).

**Table 2.** Variant call summary data.

| Data | No filters | QUAL and DP filters | Final Filters |
| --- | --- | --- | --- |
| Total number of sites | 8.031.682 | 7.960.684 | 37.027 |
| Variants | 75.607 | 55.402 | 37.027 |
| SNVs | 69.227 | 51.230 | 35.361 |
| INDELS | 6.380 | 4.172 | 1.666 |
| Quality* Mean | 269.695 | 271.213 | 283.612 |
| DP** Mean | 283.832 | 226.061 | 279.961 |

(\*) Phred Score; (\*\*) Depth of coverage.

Regarding the number of 55,402 SNVs, it can be said that it is relatively close to what was found in the literature, citing the work done by Vanburen *et al*. in 2015, where they obtained 66,579 SNVs using similar filtering metrics and the same samples used in the present study. However, Ueno *et al*. in 2014 reported 14,528 SNVs when comparing HSY and MSY mapped against the HSY reference genome. This study mentions an average of 1.8 SNVs per kbp (kilobase pairs), with the present study finding 8.62 SNVs per kbp when considering the total number of variants, and 4.4 SNVs per kbp when counting only those that passed the filters. It is worth noting that the process of identifying SNVs in cases of sexual determination has methodological support from the literature, such as the study by Jeffries, Mee, and Peichel (2012), which identified a candidate gene for sexual determination in the fish species *Culaea inconstans* from studies with variants.

### 3.2 Association Analyses

At this stage, 37,027 SNVs were subjected to analysis using PLINK. The analysis revealed that 14,982 SNVs had a p-value less than 0.05 and 8,289 SNVs had a p-value below 0.01. The number of significant variants is consistent with findings in the literature, notably mentioning Ueno *et al*. (2014), who identified 14,528 SNVs, although the authors did not distinguish how many of these were significant or if all were significant. Other researchers who conducted studies with SNVs differing between MSY and HSY include Liao, Yu, and Ming (2017), who found 21,088 SNPs and 1,887 indels. In other association studies, Geraldes *et al*. (2015) identified 650 significant SNVs in other plant species, where 13 sex-associated genes were identified after performing a genome-wide association study (GWAS).

The Manhattan plot (Figure 4) allows visualization of data from the variant calling analyzed by PLINK to investigate SNVs associated with sexual determination between males and hermaphrodites. It displays the statistical significance of each variant, facilitating the identification of SNVs potentially associated with sexual determination. Columns of clustered points, such as those found at the right end of the graph, suggest an area of interest to be investigated further, searching for mutations in genes that are indeed associated with sexual determination.

**Figure 4.**
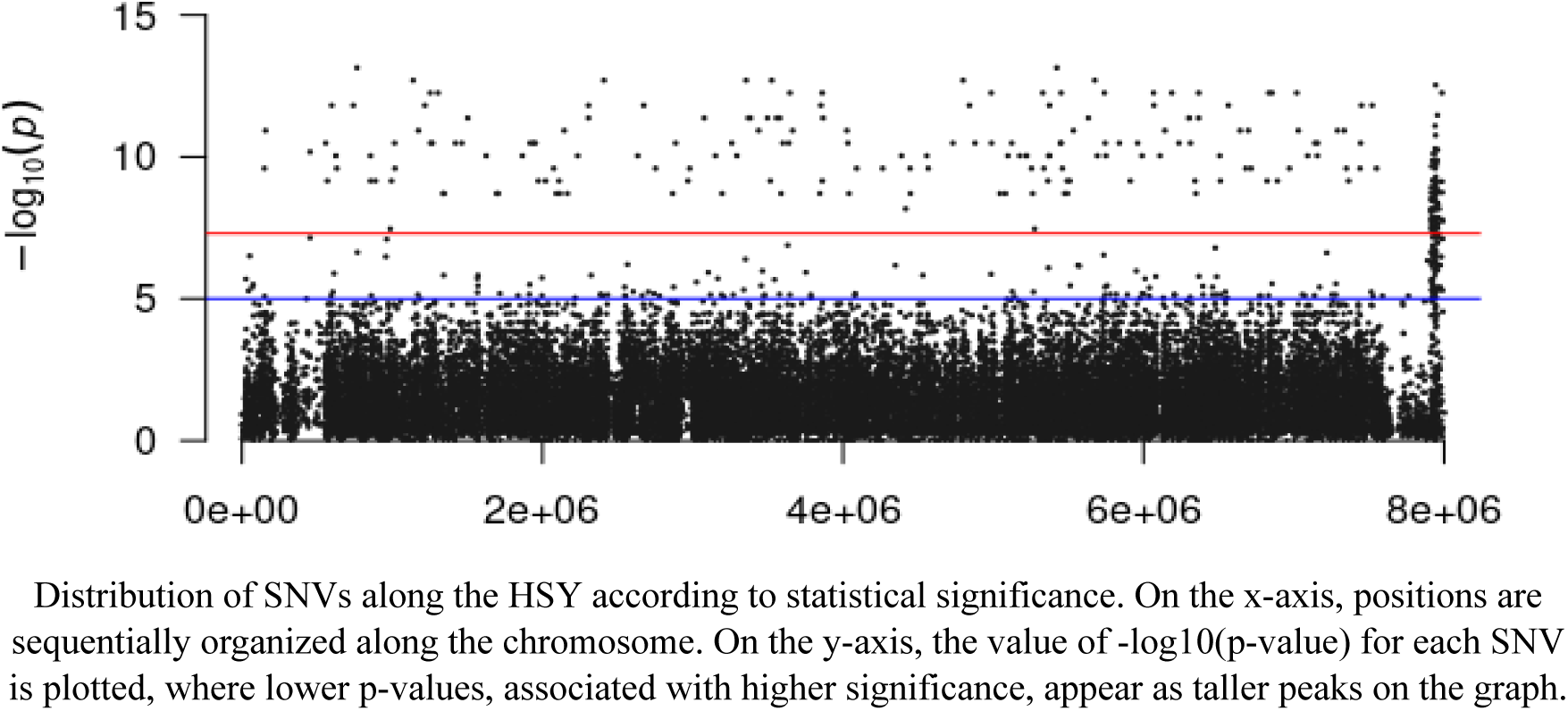
Manhattan Plot showing the distribution of SNVs according to chromosome position and their respective p-values on a logarithmic scale.

From the variants present in the Manhattan plot, 251 variants were selected that were above the red horizontal line. This line represents the genome-wide significance threshold where P = 5 x 10^-8. These variants were chosen for demonstrating a high level of significance, as they mostly represent sites where all samples of one phenotype possessed a specific nucleotide, unlike the group with the other phenotype, or vice versa. Therefore, it became necessary to evaluate the variants individually by predicting their effects using the SnpEff software.

It is important to note that even though these variants are statistically more likely to impact sexual determination in *C. papaya*, their association is not guaranteed. Other factors such as sample size and quantity must be considered, as well as factors like *linkage disequilibrium*, which can be defined as a non-random association of alleles at different loci in a population (Slatkin, 2008). One way to obtain statistics that mitigate the possibility of false positives is by performing a Bonferroni adjustment, which the PLINK output files also provide (Bonferroni, 1936). The Bonferroni method is a statistical technique used to control Type I error (false positives) in multiple hypothesis testing. It is frequently applied in genome-wide association studies (Gao *et al*., 2010). With the Bonferroni adjustment, 309 significant variants were counted in the present study.

### 3.3 SnpEff

Among the results obtained from SnpEff for the prediction of the effects of the 251 variants (Table 4), it became evident that mutations constituting transitions were more common than transversions (Table 5), which was expected. This is because changes between purines and pyrimidines, as seen with transversions, require larger chemical changes in the molecular structure. Consequently, these mutations tend to be more detrimental to protein function and, therefore, fewer transversions persist through natural selection (Lyons & Lauring, 2017).

**Table 4.** Variantes applied to SnpEff.

| Variant Type | Quantity |
| --- | --- |
| SNVs | 242 |
| Deletions | 2 |
| Insertions | 7 |

**Table 5.**
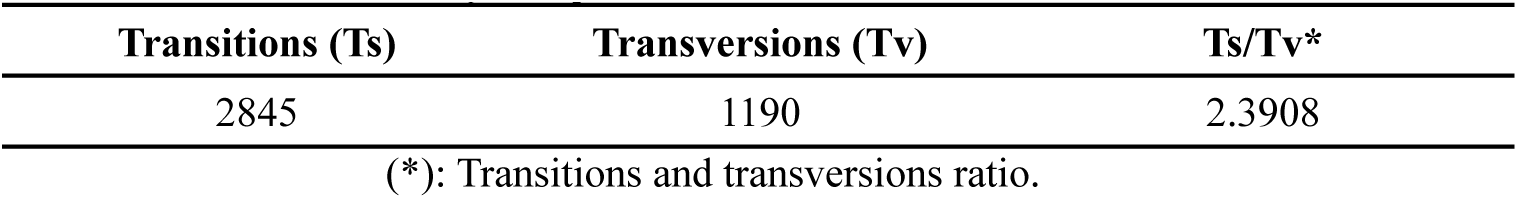
Quantity of transitions and transversions.

| Transitions (Ts) | Transversions (Tv) | Ts/Tv* |
| --- | --- | --- |
| 2845 | 1190 | 2.3908 |

(\*): Transitions and transversions ratio.

In total, SnpEff predicted 449 effects (Table 6), with the majority being of moderate impact and none of high impact. It is also noteworthy that most of the predicted effects are not located in coding regions (Table 7). These results are consistent with similar studies, such as Bohry *et al*. (2021), who found only 4% of their effects in coding regions. Regarding the degree of effects, it is important to conceptualize that predicted effects are categorized into four types: (1) high represents effects from nucleotide changes that can disrupt translation, alter the reading frame, and potentially generate truncated proteins with loss of function; (2) moderate reflects effects of a non-synonymous mutation in a nucleotide, meaning it produces a different amino acid during translation, which can directly affect protein function and structure; (3) low are effects from synonymous mutations that do not cause amino acid changes; (4) modifier includes effects from mutations occurring in non-coding regions (Cingolani *et al*., 2012). The quantity of effects, their impact degrees, effect types, and their locations can be seen in the tables below:

**Table 6.** Predicted Effects by SnpEff.

| Impact Level | Quantity |
| --- | --- |
| Modifier<sup>1</sup> | 402 |
| Low<sup>2</sup> | 22 |
| Moderate<sup>3</sup> | 25 |
| High<sup>4</sup> | 0 |

(1) Modifier: effects of mutations occurring in non-coding regions; (2) Low: effects of synonymous mutations; (3) Moderate: effects of non-synonymous mutations; (4) High: effects of nucleotide changes that can disrupt translation, alter the reading frame, or generate truncated proteins with loss of function.

**Table 7.** Distribution of predicted effects in SnpEff according to categories.

| Category | Quantity |
| --- | --- |
| 3_prime_UTR<sup>1</sup> | 11 |
| 5_prime_UTR<sup>2</sup> | 4 |
| Downstream Gene<sup>3</sup> | 106 |
| Intergenic<sup>4</sup> | 129 |
| Intron<sup>5</sup> | 61 |
| Missense<sup>6</sup> | 25 |
| Splice Region<sup>7</sup> | 6 |
| Synonymous<sup>8</sup> | 16 |
| Upstream Gene<sup>9</sup> | 91 |
| Exon<sup>10</sup> | 40 |

(1) Variant in the 3' untranslated region of the gene. (2) Variant in the 5' untranslated region of the gene. (3) Variant downstream of the gene. (4) Variant in intergenic regions. (5) Variants in introns. (6) Variant that alters the codon and the amino acid. (7) Variant in the splicing region of pre-mRNA. (8) Variant that alters the codon but not the amino acid. (9) Variant upstream of the gene. (10) Variant in the coding region.

Therefore, this dissertation aimed to investigate the 25 moderate effects identified (Table 8) by SnpEff. To do this, BLASTPs were performed with the genes exhibiting these effects, predicted by the Augustus software, using their respective amino acid sequences. This measure was taken to determine whether these genes are somehow related to the candidate genes for sexual determination in *C. papaya* previously mentioned.

**Table 8.** Information about the moderate effects predicted by SnpEff.

| Position | QUAL SCORE | REF | ALT | P-VALUE | Gene | Type | Amino acid |
| --- | --- | --- | --- | --- | --- | --- | --- |
| 1018426 | 285 | A | G | 2.54e-10 | g84 | MISSENSE | Met308Val |
| 1217284 | 285 | A | C | 1.537e-128 | g106 | MISSENSE | Phe57Val |
| 1502921 | 285 | T | C | 4.262e-129 | g136 | MISSENSE | Arg67Gly |
| 2409162 | 285 | T | A | 2.005e-13 | g238 | MISSENSE | Asp135Glu |
| 3557335 | 285 | C | G | 4.262e-12 | g329 | MISSENSE | Leu101Val |
| 3579477 | 285 | C | G | 4.262e-12 | g331 | MISSENSE | Pro67Gln |
| 4090833 | 284 | A | G | 3,19E-08 | g392 | MISSENSE | Arg46Gly |
| 4264109 | 285 | A | G | 2.54e-10 | g410 | MISSENSE | Ile59Val |
| 4799473 | 285 | T | C | 2.005e-13 | g467 | MISSENSE | Ile334Thr |
| 5217967 | 278 | A | G | 9.127e-11 | g504 | MISSENSE | Met100Thr |
| 6834317 | 285 | T | C | 5.55e-13 | g642 | MISSENSE | Leu349Pro |
| 7030337 | 285 | A | G | 1.183e-11 | g658 | MISSENSE | Gln131Arg |
| 7914676 | 285 | A | G | 1.822e-09 | g746 | MISSENSE | Thr76Ala |
| 7915788 | 285 | G | T | 3.102e-09 | g746 | MISSENSE | Lys311Asn |
| 7916380 | 285 | A | C | 7.869e-09 | g746 | MISSENSE | Asn449His |
| 7916829 | 285 | A | G | 2.156e-10 | g746 | MISSENSE | Asn575Asp |
| 7940417 | 285 | T | G | 8.081e-12 | g747 | MISSENSE | His1059Pro |
| 7940763 | 285 | A | G | 9.141e-10 | g747 | MISSENSE | Ile973Thr |
| 7941853 | 285 | C | T | 6.769e-09 | g747 | MISSENSE | Ala778Thr |
| 7942156 | 285 | T | C | 4.011e-09 | g747 | MISSENSE | Ile712Val |
| 7942162 | 285 | T | C | 2.024e-08 | g747 | MISSENSE | Lys710Glu |
| 7942336 | 285 | C | T | 1.822e-09 | g747 | MISSENSE | Val652Ile |
| 7944092 | 285 | C | G | 1.511e-09 | g747 | MISSENSE | Ser348Thr |
| 7944291 | 285 | C | T | 6.267e-09 | g747 | MISSENSE | Val282Ile |
| 7945174 | 285 | T | A | 4.628e-08 | g747 | MISSENSE | Tyr184Phe |

(1) Position, variant nucleotide site between chromosomes Y and Yh; (2) Qual Score, alignment quality score between the sequences and the HSY reference genome; (3) (4) REF and ALT, reference and alternative nucleotides, respectively; (5) P-value, statistical value calculated by PLINK; (6) Gene predicted in Augustus (default names); (7) Type, characteristic of the SNV type; (8) Reference amino acid, its location in the protein sequence, and the alternative amino acid.

### 3.4 BLASTP

The results of the BLASTp with the sequences of genes with moderate effects, performed through the NCBI platform, included many results with retrotransposons, some polymerases, gag-pol proteins, integrases, reverse transcriptases, nucleotidyltransferases, ribonucleases, endonucleases, and some chromatin factors. Many sequences from other plants such as grape (*Vitis sp.*), tangerine (*Citrus reticulata*), cotton (*Gossypium hirsutum*), coffee (*Coffea arabica*), wild potato (*Solanum jamesii*), and notably the similarities found with melon (*Cucumis melo*), which is also under investigation to uncover its sexual determination process, were found.

Finally, it is relevant to reiterate that some of these genes (from Augustus prediction) did not return sequences with similarity, and those that did had approximately 80-90% of the query sequence covered in alignment with the sequences found by BLASTp. The high number of mobile element sequences is noteworthy, considering how these elements may have evolved in relation to the *C. papaya* genome, specifically concerning the recombination between Y and Yh. Therefore, it is essential to further analyze and delve into these results in future studies.

### 3.5 De novo genome assembly of the male papaya sample and alignment

After the scaffolds were assembled using SPAdes and aligned with the genome of a hermaphrodite sample to investigate the differences, two distinct graphs were generated. The first (Figure 5) is the alignment of all male scaffolds with the entire hermaphrodite genome. The second (Figure 6) is an alignment of only the scaffold belonging to the MSY region with the HSY region. Common to both dot plots generated in D-genies is a continuous ascending line, suggesting a good overall match between the compared genomes. However, there are also some small gaps, and in Figure 5, a notable discrepancy in size between the scaffold set (377 Mb) and the complete hermaphrodite reference genome (339 Mb) is observed.

**Figure 5.**
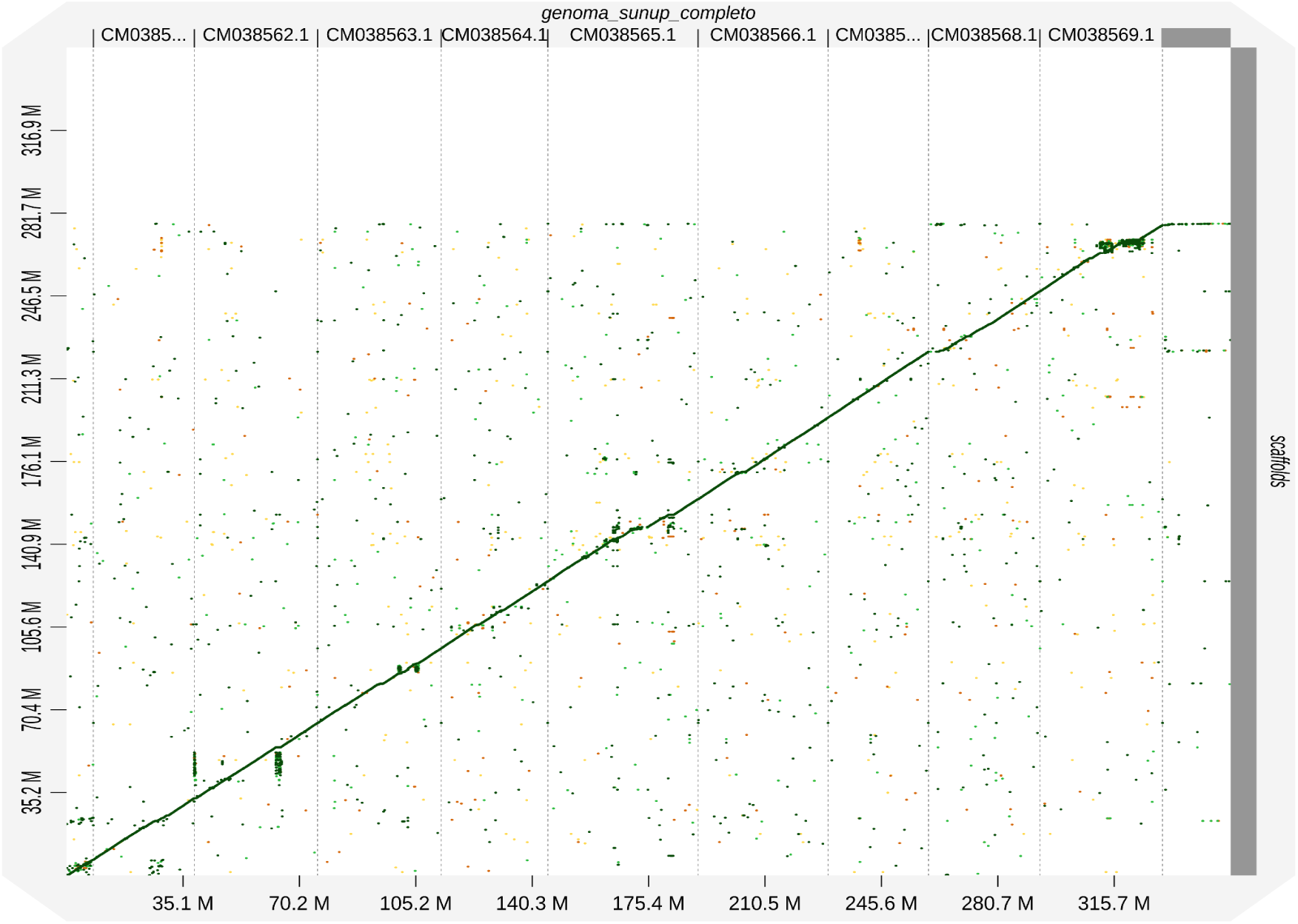
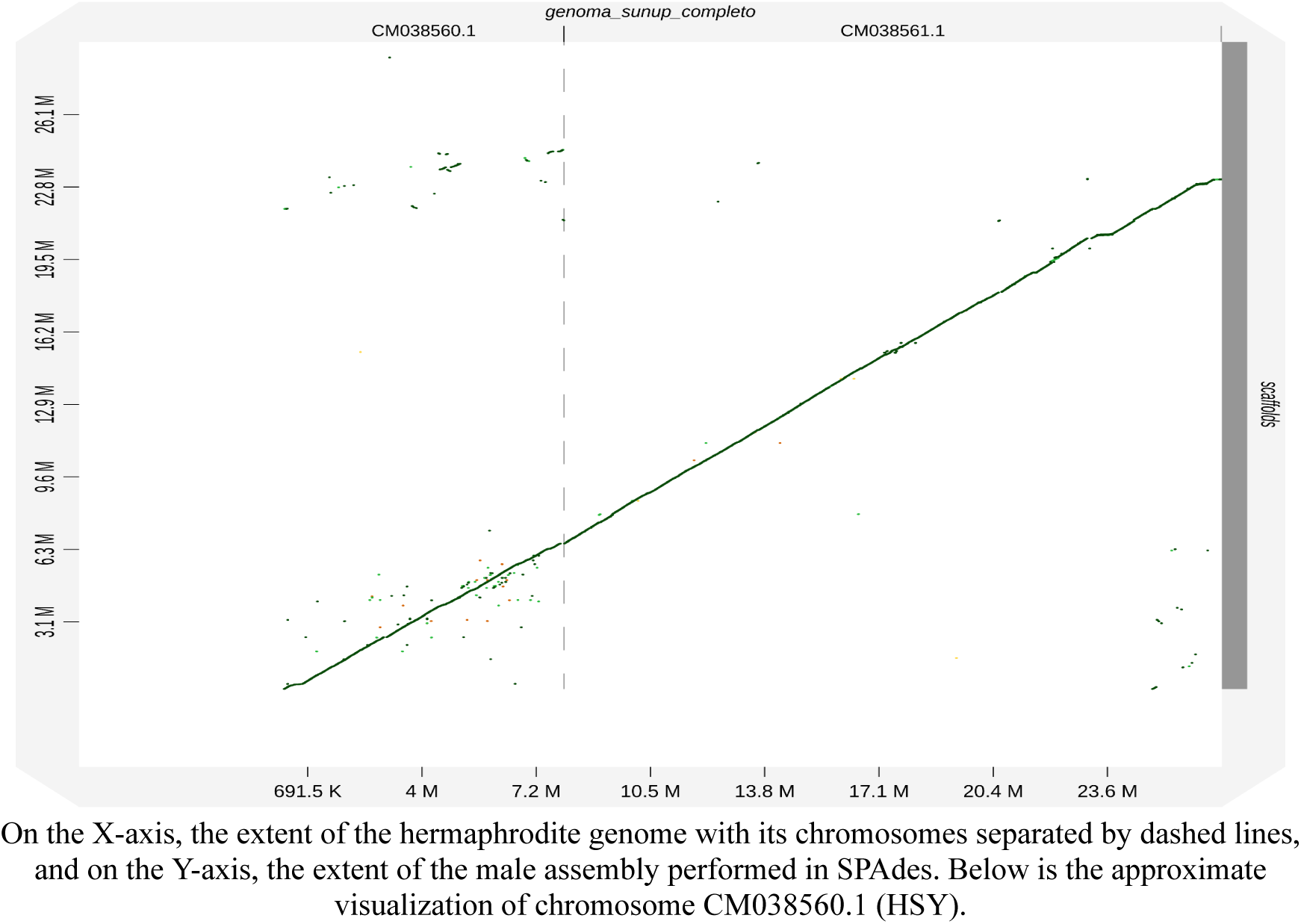
*Dot plot* of the complete hermaphrodite genome aligned with the scaffolds created by SPAdes.

**Figure 6.**
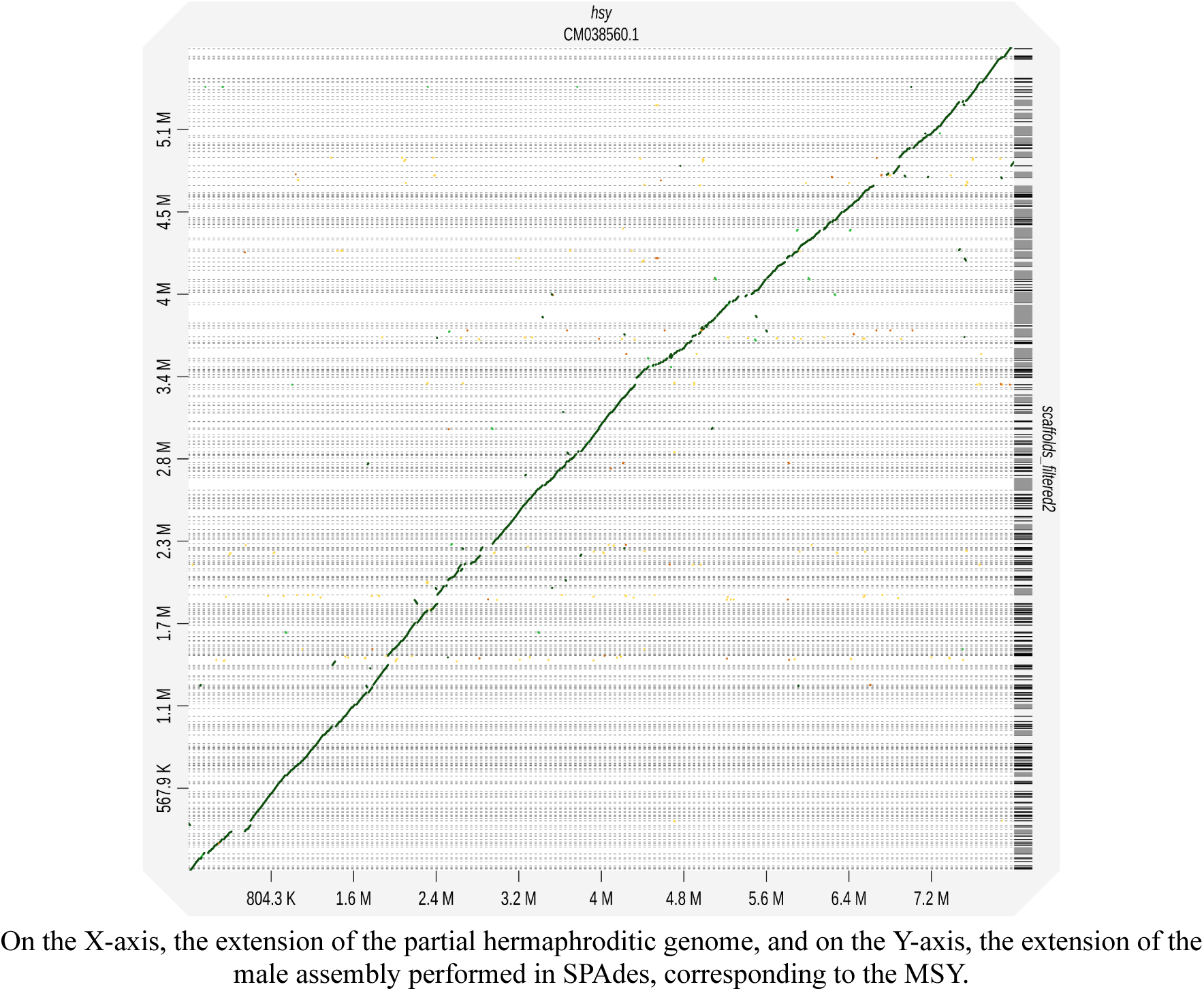
Dot plot of the alignment between the HSY and the male scaffold that corresponds to the MSY.

## 4. Conclusion

The present work provides promising data but still requires further in-depth analysis. However, some information becomes evident, such as the presence of SNVs associated with differentiation between male and hermaphrodite phenotypes, and the predicted effects by SnpEff may have biological significance. Future perspectives include conducting more detailed computational analyses to provide greater insight into the influence of these SNVs on sexual determination in *C. papaya*. It is anticipated that bench experiments could be employed to evaluate the impact of these mutations on the manifestation of sexual phenotypes in vivo. Additionally, it is important to conduct similar studies comparing the differentiation between female papayas and male/hermaphrodite papayas. It is hoped that this research can serve as a starting point for a deeper understanding of sexual determination, not only in *C. papaya* but also in species with similar sexual segregation during the initial process of chromosome differentiation.

## Notes

### Competing Interest Statement

The authors have declared no competing interest.

